# Interaction between Discs large and GPSM2: A Comparison Across Species

**DOI:** 10.1101/2021.07.31.454595

**Authors:** Emily Schiller, Dan T. Bergstralh

## Abstract

The orientation of the mitotic spindle determines the direction of cell division, and therefore contributes to tissue shape and cell fate. Interaction between the multifunctional scaffolding protein Discs large (Dlg) and the canonical spindle orienting factor GPSM2 (also called Pins in *Drosophila* and LGN in vertebrates) has been established in bilaterian models, but its function remains unclear. We used a phylogenetic approach to test whether the interaction is obligate in animals, and in particular whether GPSM2 evolved in multicellular organisms as a Dlg-binding protein. We show that Dlg diverged in *C. elegans* and the syncytial sponge *O. minuta* and propose that this divergence may correspond to differences in spindle orientation requirements between these organisms and the canonical pathways described in bilaterians. We also demonstrate that GPSM2 is present in basal animals, but the established Dlg-interaction site cannot be found in either Placozoa or Porifera. Our results suggest that the interaction between GPSM2 and Dlg appeared in Cnidaria, and we therefore speculate that it may have evolved to promote accurate division orientation in the nervous system. This work reveals the evolutionary history of the GPSM2/Dlg interaction and suggests new possibilities for its importance in spindle orientation during epithelial and neural tissue development.

## Introduction

Discs large (Dlg) is a multifunctional scaffolding protein (Noirot-Timothée & Noirot, 1980; Woods et al., 1996; Woods & Bryant, 1994). Like other MAGUK (membrane associated guanylate kinase) proteins, Dlg is comprised of three N-terminal PDZ domains, an SH3 domain, and a C-terminal GUK (Guanylate Kinase) domain that is catalytically inactive and binds phosphorylated proteins (Woods et al., 1996; Woods & Bryant, 1994; Zhu, Shang, et al., 2011; Zhu et al., 2012). Though well-studied for its roles in epithelial cell polarity and synaptic function, Dlg has also been identified as a factor involved in orientation of the mitotic spindle (Bellaïche et al., 2001; Bergstralh et al., 2013; Chanet et al., 2017; Johnston et al., 2009; Nakajima et al., 2013; Saadaoui et al., 2014; Siegrist & Doe, 2005)

The canonical spindle orientation complex is comprised of at least three evolutionarily conserved proteins: NuMA (nuclear mitotic apparatus), GPSM2 (G-protein signaling modulator, also called LGN), and the microtubule motor protein dynein (Figure 1). NuMA and dynein combine to produce a pulling force that reels the spindle into alignment by walking towards the minus end of astral microtubules. GPSM2 is thought to work as an adaptor; it binds to NuMA through its N-terminal TPRs (tetratricopeptide repeats) and to myristoylated Gαi through its C-terminal GoLoco domains, thereby anchoring the complex to the cell cortex (Figure 1). This structure is conserved across taxa, though the number of TPR and GoLoco domains varies between and within phyla (Wavreil & Yajima, 2020) A notable exception is the nematode worm, in which two functional homologs to GPSM2, called GPR1 and 2, have a unique structure that lacks GoLoco domains and may contain one or two TPRs, though this is debated (Nguyen-Ngoc et al., 2007; Siller & Doe, 2009; Willard et al., 2004).

**Figure 1:**
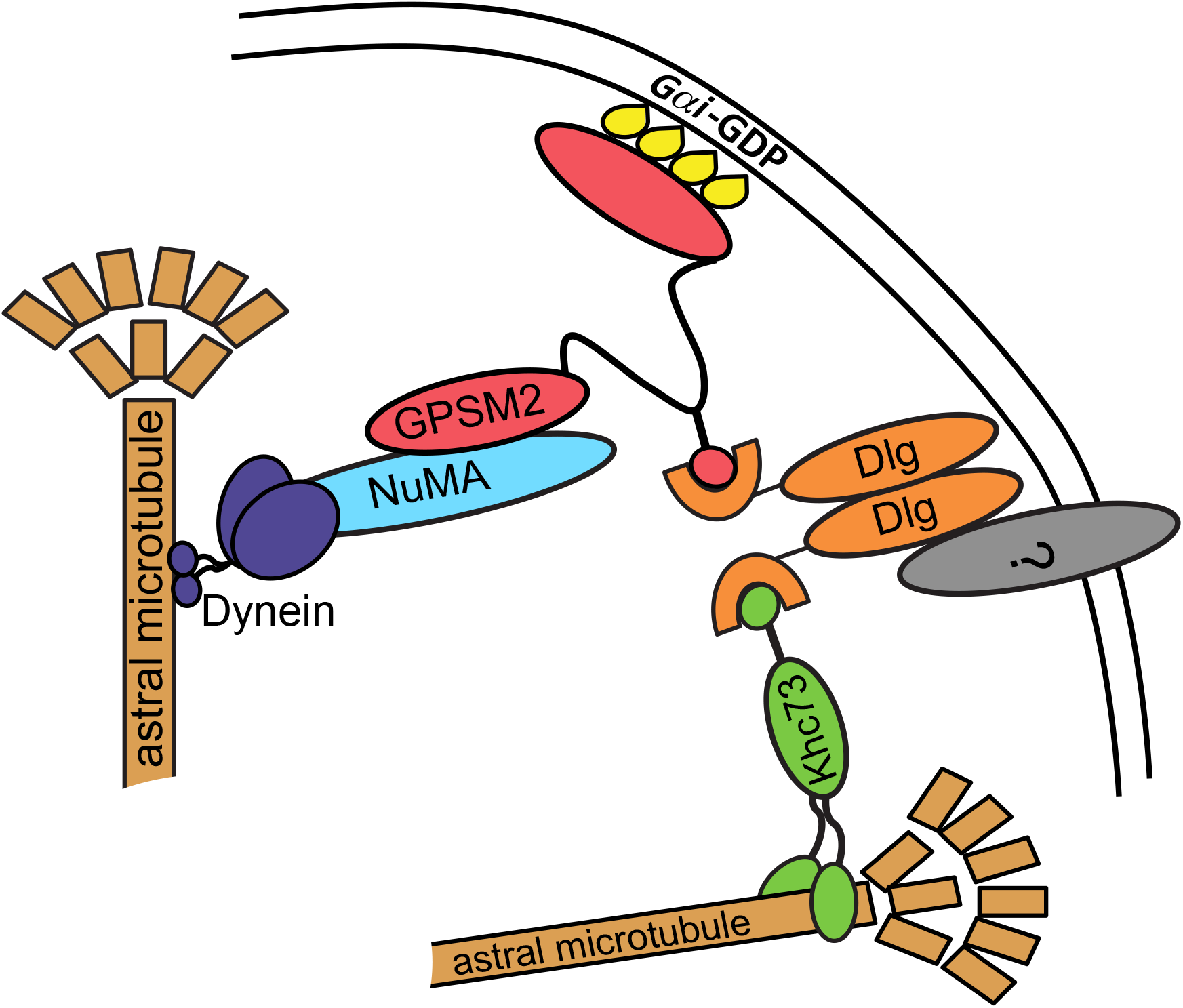
The Canonical Spindle Orientation Complex. The spindle orientation complex proteins NuMA (light blue), GPSM2 (fuchsia), and Dlg (orange). Khc73 (pink), a potential spindle orientation factor, is also shown. This complex functions to capture microtubules and link the motor protein dynein (dark blue) to the cortex.

Extensive evidence links Dlg to spindle orientation through direct interaction with GPMS2, mediated by the Dlg GUK domain (Bellaïche et al., 2001; Johnston et al., 2012) Crystallization of GPSM2/Pins and Dlg from *D. melanogaster* and *M. musculus* reveals that binding relies on phosphorylation of GPSM2 at a serine residue (S436 in *D. melanogaster,* S401 in vertebrates) in the unstructured linker between the TPR and GoLoco domains (Johnston et al., 2012; Woods et al., 1996; Zhu, Shang, et al., 2011). In agreement, expression of the non-phosphorylatable S401A variant in MDCK cell cysts or chick neuroepithelium causes spindle misorientation at metaphase (Hao et al., 2010; Saadaoui et al., 2014). Additionally, *Drosophila* follicle epithelial cells homozygous for *dlg^1P20^*, which encodes a premature truncation that disrupts the GUK domain, orient spindles randomly (Bergstralh et al., 2013).

Two models, which are not exclusive, have been proposed to explain the function of this interaction. The first is that Dlg functions as a positioning cue that recruits and/or restricts GPSM2 to distinct regions of the cell cortex (Bergstralh et al., 2013; Saadaoui et al., 2014). The second model is that Dlg acts to link the canonical complex to a microtubule motor called Khc73, thereby promoting microtubule capture (Siegrist & Doe, 2005; Zhu et al., 2016). This interaction also utilizes the GUK domain, which can bind the Khc73 MBS (MAGUK binding stalk) domain in a manner that relies on phosphorylation at a region outside the binding site (Zhu et al., 2016). A challenge faced in reconciling these two models is the problem of how Dlg could interact with both proteins in the same spindle orientation complex. However, *in vitro* studies suggest that Dlg exists as a dimer, in which case one monomer could bind to GPSM2 and the other to Khc73 (Fig. 1) (Marfatia et al., 2000).

### Spindle orientation requirements for neural tissue

In mammals, the phenotypic changes associated with spindle misorientation are predominantly observed in the nervous system, where they underlie diseases such as primary microcephaly, Chudley-McCullough syndrome, and Huntington’s disease (Barnat et al., 2020; Godin et al., 2010; Godin & Humbert, 2011; Higgins et al., 2010; MacDonald et al., 1993; Noatynska et al., 2012). For example, exome sequencing and homozygosity mapping identified a mutation in human GPSM2 associated with the sensorineural hearing loss and brain malformations in Chudley-McCullough syndrome (Doherty et al., 2012) Additional experiments in mice demonstrated that this mutation causes profound deafness in otherwise phenotypically normal mice (Bhonker et al., 2016). Since the truncated GPSM2 mutant lacks GoLoco domains, the interaction between GPSM2 and Gαi is disrupted, but only appears to affect hearing and brain formation. Indeed, Gαi is essential for central auditory processing, inner hair cell synapse maturation, and hair bundle shape in mice (Beer-Hammer et al., 2018).

Neural defects due to spindle misorientation are also observed outside of vertebrates. In *Drosophila, mud* (NuMA) mutants are viable and demonstrate only a few defects: morphological changes in the mushroom bodies of the brain (Bowman et al., 2006; Prokop & Technau, 1994), slightly smaller wings, and female sterility (Zhou et al., 2019).

Variations in the requirements for spindle orientation between different tissues is well established. An important example is the presence of the protein Insc (Inscuteable) in neuroblasts but not in epithelia. In bilaterian models, Insc orients asymmetrical neuroblast division in the apical-basal axis by apically localizing GPSM2 (Yu et al., 2000). Unlike Dlg which interacts with GPSM2 at the phosphorylated linker, Insc localizes GPSM2 by binding to the TPR domains (Yuzawa et al., 2011). Interestingly, the asymmetric localization of GPSM2 in neuroblast spindle orientation can be mediated through either Insc or Dlg, each with a distinct pathway (Siegrist & Doe, 2005).

While the importance of the interaction between Dlg and GPSM2 has been demonstrated in bilaterian models, whether these proteins and their interactions are conserved in more ancient animals remains unknown. If this interaction is conserved across all animal phyla, then the interaction between GPSM2 and Dlg may be essential to multicellularity. Alternatively, the interaction could have evolved as tissue types and architectures diversified within the animal kingdom. Therefore, determining when the Dlg and GPSM2 interaction arose may reveal the possible functions of the interaction in spindle orientation.

## Results

### The GUK domain is conserved in basal animals

The Dlg GUK domain is found in animals and the single-celled eukaryotes choanoflagellates and Filasterea (Mendoza et al., 2010; Velthuis et al., 2007). Our analysis indicates that the GUK domain is evolutionarily conserved; global alignment reveals sequence similarity at the Dlg C-terminus, and computationally predicted structures at this region resemble the experimentally determined structure (Figures 2A,B). Conserved residues at the Dlg GUK domain, specifically at the GB (GMP binding) subdomain, that interact with the phosphorylated GPSM2 linker have been identified in *H. sapiens, M. musculus, D. rerio,* and *D. melanogaster* (Anderson et al., 2016; Johnston et al., 2011, 2012). Included in these residues is the proline residue responsible for the functional switch from GMP to phosphoprotein binding activity. We extended this analysis to other bilaterians and found that the GB subdomain is highly conserved. Similarity is also evident outside of bilaterian animals, with on average 15% greater identity locally than globally in Cnidaria, Placozoa, and Porifera, though we could not identify a Dlg homolog in Ctenophora (Figure 2C). We also identified Dlg homologs in the choanoflagellates *S. rosetta* and *O. brevicollis* and show here that GB domains in these proteins share an average of 54% and 56% identity (respectively) with animals (Figure 2C). Together, these results agree with previous work suggesting that the ability of Dlg to bind GPSM2 has been retained throughout its evolutionary history (Anderson et al., 2016). We also identified two notable exceptions; *C. elegans* and the poriferan *Oopsacas minuta.* For both organisms, we observed sequence divergence from other animals in the global and GB domain alignments (Figure 2C and Supplemental Figure 1A,B).

**Figure 2:**
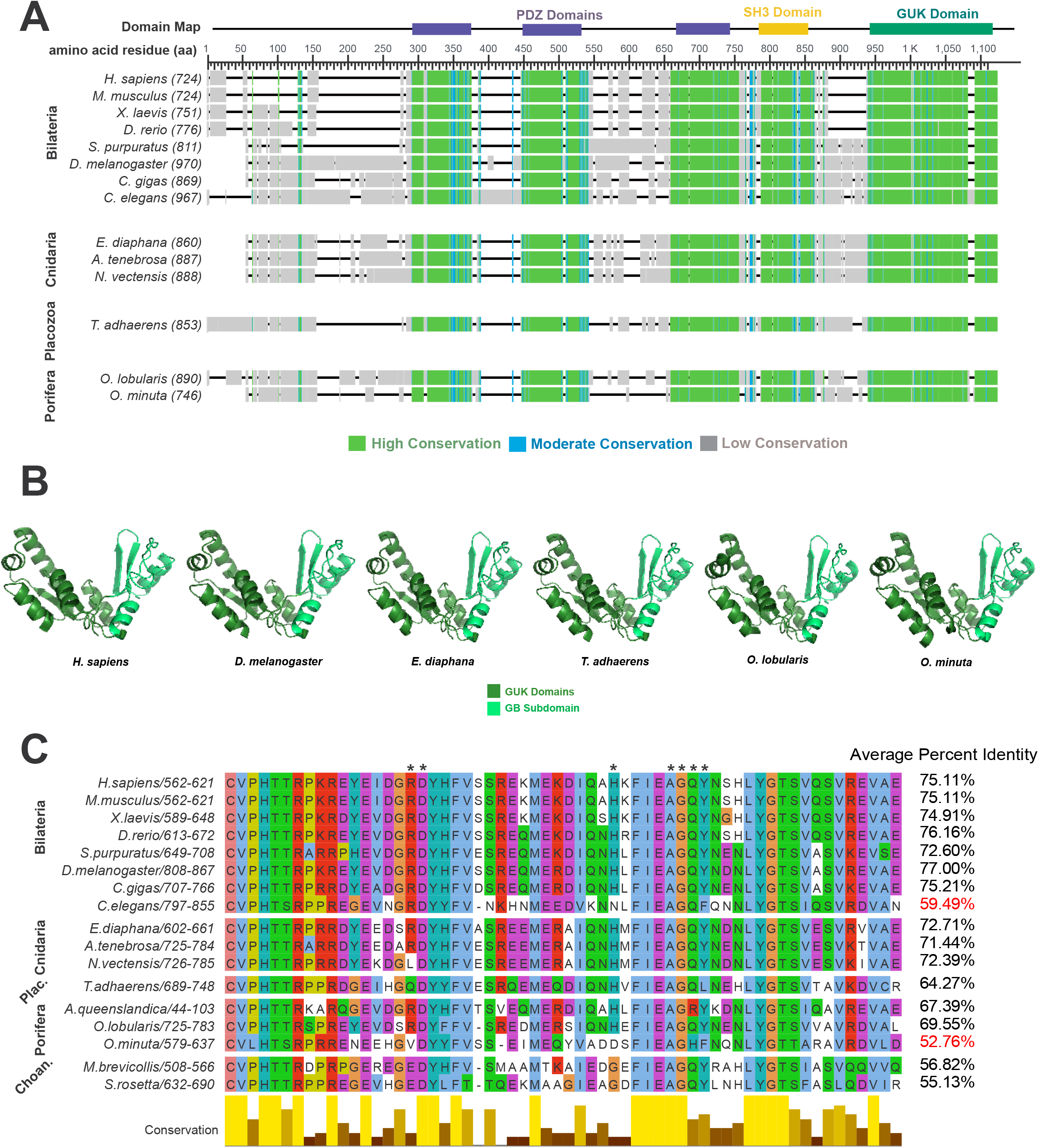
Evolutionary Conservation of Discs large. A) Dlg sequences are aligned and colored based on conservation. The *H. sapiens* Dlg domain map is aligned to conservation map. Figure is adapted from the output of the Multiple Sequence Alignment Viewer application (Papadopoulos & Agarwala, 2007). B) Predicted structure of Dlg. Models were created using the trRosetta in the Robetta server and edited in PyMOL Molecular Graphics System, Version 2.4.1. C) Dlg GB subdomain sequence alignment with organisms included in this study and choanoflagellates (M. brevicollis & S. rosetta). Asterisks indicate residues that are important for the interaction with phosphorylated GPSM2. Alignment is colored using the ClustalX color scheme. Figure is adapted from Jalview 2 (Waterhouse et al., 2009).

### Syncytial sponges lack conservation at the GUK-GB subdomain

There are four classes of sponge within the Porifera phylum: Calcarea, Demospongia Hexactinellida, and Homoscleromorpha. Hexactinellida cell types are mainly syncytial, while Calcarea, Demospongia, and Homoscleromorpha contain entirely mononuclear cells. We found that the Dlg protein sequence has diverged in *O. minuta,* a member of Hexactinellida. Of the seven amino acids that interact with the GPSM2 linker, only three are conserved in *O. minuta* while five are conserved in *A. queenslandica* (Demospongiae) and all seven are conserved in *O. lobularis* (Homoscleromorpha). Furthermore, the functional proline residue is conserved in *A. queenslandica* and *O. lobularis* but is replaced by a leucine in *O. minuta* (Fig. 2C) (Johnston et al., 2012). We considered the possibility that these differences could reflect low quality of the *O. minuta* genome annotation. To test for this, we examined the polarity proteins aPKC and Scribble. These sequences are equally conserved across all three sponges (not shown).

### The Khc73 MBS domain is conserved across animals

Before investigating conservation of the Dlg-interacting region in GPSM2, we first asked to what extent sequence conservation should be expected. To address this question, we examined the Dlg-interacting region in Khc-73, a well-established binding partner for Dlg, with particular attention paid to sequences in animals outside of the Bilaterian clade (Figure 3A). Khc73 is globally conserved across *H. sapiens*, the choanoflagellate *S. rosetta,* and Filasterea *C. owczarzaki* (Anderson et al., 2016) and locally conserved at a 14-3-3 binding motif across *H. sapiens, X. laevis, D. rerio,* and *D. melanogaster* (Lu & Prehoda, 2013). However, it is unknown whether the MAGUK-binding stalk is conserved within the animal kingdom. We found that the MBS domain sequence is highly conserved across phyla (Figure 3B and Supplemental Figure 1C).

**Figure 3.**
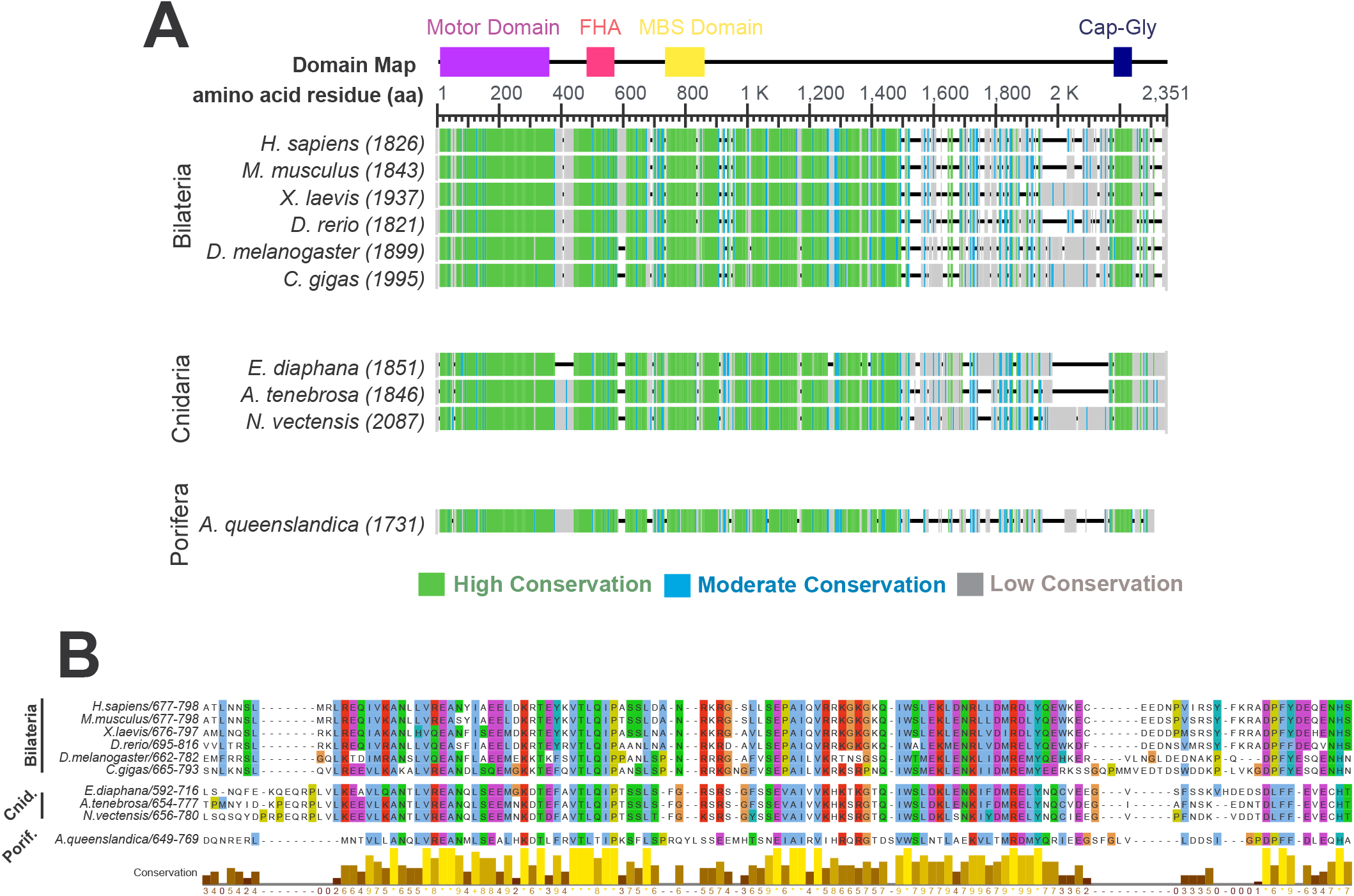
The Khc73 MBS domain is locally conserved in Porifera, Cnidaria, and Bilateria. A) Khc73 sequences are aligned and colored based on conservation. The *H. sapiens* Dlg domain map is aligned to conservation map. B) Sequence alignment of identified Khc73 MBS domains. Alignment is colored using the ClustalX color scheme.

### Identification of a candidate GPSM2 in Porifera

We next sought to identify GPSM2 orthologs in basal animals. BLAST searching against Ctenophora and Porifera genomes recovered no hits for GPSM2 in ctenophores and only one hit, a GPSM2-like protein (XP_019856151.1; NW_003546593.1 (14479…19908, complement)) in the sponge *A. queenslandica.* This sequence lacked an apparent linker region and GoLoco domains. Manual analysis of the *Amphimedon queenslandica* Annotation Release 102 genomic sequence revealed an additional annotated sequence with homology to the GoLoco domains (XP_003389018.1; NW_003546593.1 (9550…10710, complement)) located 3,769 nucleotides past the C-terminus of XP_019856151.1. These sequences were combined to generate a candidate full length GPSM2 sequence (Figure 4A and Supplemental Figure 1D).

**Figure 4.**
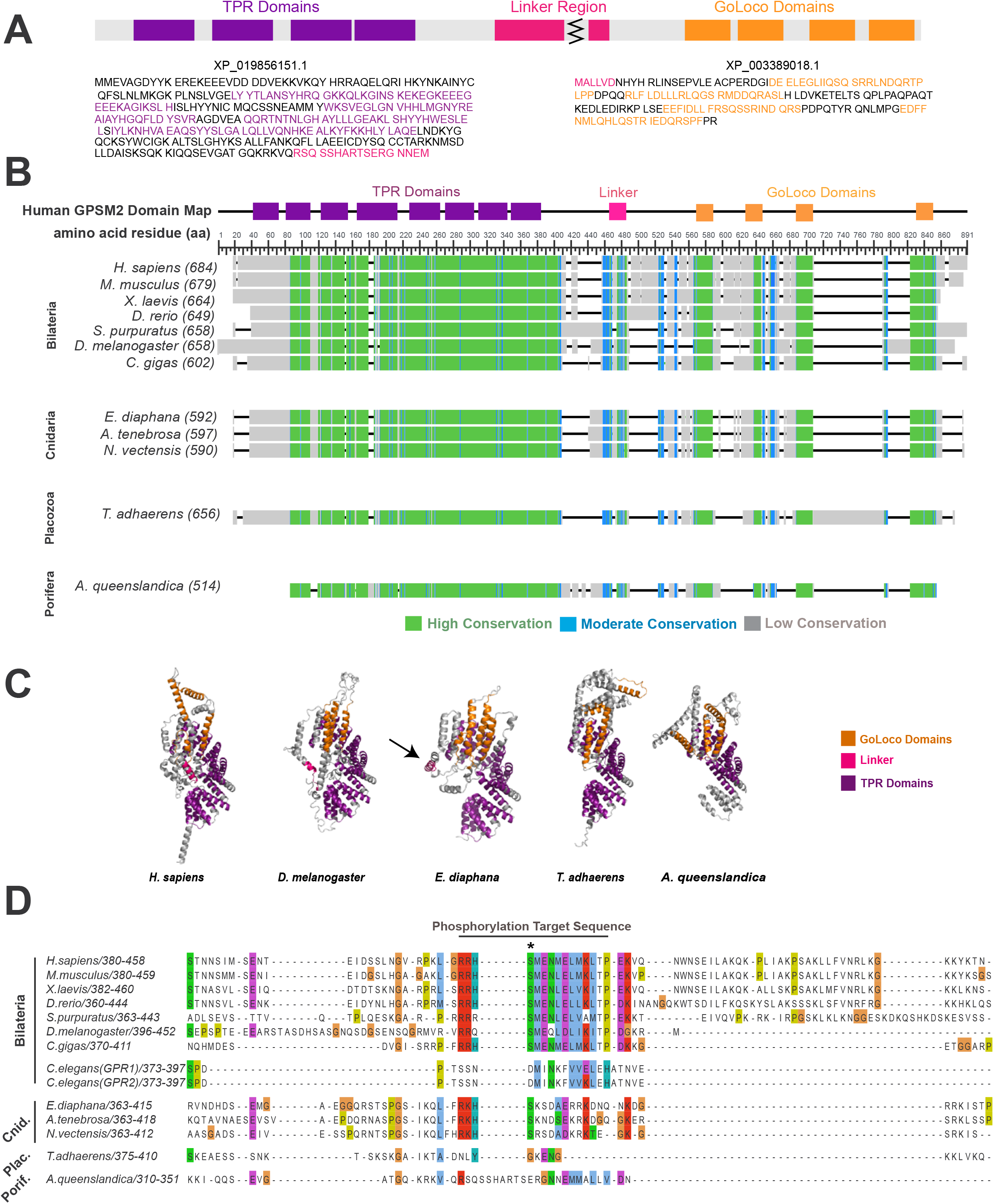
Evolutionary Conservation of GPSM2. A) A reconstructed sequence for the GPSM2 ortholog in the poriferan *A. queenslandica.* B) Identified GPSM2 sequences are aligned and colored based on conservation. The H. sapiens GPSM2 domain map is aligned to conservation map. Figure is adapted from the output of the Multiple Sequence Alignment Viewer application (Papadopoulos & Agarwala, 2007). C) Predicted structure of GPSM2. Models were created using trRosetta as in Figure 2. D) Sequence alignment at the GPSM2 linker region. (+) in consensus sequences indicate positions where two residues share the modal value. Alignment is colored with ClustalX color scheme. Figure is adapted from Jalview 2 (Waterhouse et al., 2009).

We considered the possibility that exons past the N-terminus of XP_019856151.1 or past the C-terminus of XP_003389018.1 were missed in the manual reconstruction. We therefore performed global sequence alignment comparing the GPSM2 sequence in *A. queenslandica* to all other organisms included in this study. Based on this analysis, missing exons are likely to be located past the N-terminus of XP_019856151.1, as all organisms except *A. queenslandica* contain an additional 20-80 amino acids at the N-terminus, corresponding to the first TPR domain in *H. sapiens* GPSM2. It is also possible that an exon is missing past the C-terminus of XP_003389018.1, although other organisms in this study only contain an additional 5-20 amino acids at the C-terminus, and these do not correspond with any functional domains.

### Conservation of the GPSM2 linker region

A GPSM2 ortholog has been identified in choanoflagellates but lacks the necessary phosphorylation site to interact with Dlg. Because this site has been previously identified in multiple animal phyla, the question arises of whether it arose to facilitate multicellularity. A challenge to addressing this question is that the representative cnidarian sequence (*Hydra vulgaris* XM_004209432.1) used in the earlier alignment is no longer annotated as part of the *H. vulgaris* genome and has been removed from the NBCI GenBank (Anderson et al., 2016). We therefore examined GPSM2 sequences from the cnidarians *A. tenebrosa, E. diaphana, N. vectensis, O. faveolate, P. damicornis*, and *S. pistillata* and identified a highly conserved region with a consensus sequence of RKHSN++ADKRK (+ denotes positions where two residues share the modal value) (Supplemental Figure 2). The conserved serine residue is predicted to be a phosphorylation site with >99% confidence (Blom et al., 1999). While *E. diaphana* and *N. vectensis* contain a second phosphorylation site at a serine residue two amino acids away from the conserved phosphorylation site, the bilaterian linker contains a glutamic acid residue at this same location and therefore this double phosphorylation is unlikely to affect the interaction between Dlg and GPSM2 structurally or electrostatically. Together, these results are consistent with the possibility that interaction between Dlg and phosphorylated GPSM2 extends to cnidarians.

Previous work suggests that the mitotic kinase Aurora A phosphorylates GPSM2 and is required for spindle orientation in cultured cells with induced cortical polarity (Johnston et al., 2009). Aurora A phosphorylates proteins that contain an R/K/N-R-X-S/T-B motif, where B is any hydrophobic residue except proline (Ferrari et al., 2005). This motif is readily apparent in bilaterian GPSM2 proteins; the relevant serine residue is situated within a highly conserved consensus sequence (RRHSMENMELMKLTP) (Anderson et al., 2016; Johnston et al., 2009; Saadaoui et al., 2017). An exception is the nematode worm. An earlier study raised the question of whether an aspartic acid in the *C. elegans* linker takes the place of the phosphorylated serine, acting as a phosphomimetic that permits constitutive binding to Dlg (Johnston et al., 2009). Given the low sequence conservation of the GB subdomain of *C. elegans* Dlg, we suggest this possibility is unlikely.

The conserved phosphorylation target sequence in cnidarians does not match the bilaterian target. All identified cnidarian linker sequences contain either a charged or polar residue (R/K/N) at the B position (Supplemental Figure 2). The efficiency (K_cat_/K_m_) of human Aurora A phosphorylation of synthetic peptides is reduced by 85% when the B position is occupied by an arginine versus a leucine (Ferrari et al., 2005). This observation suggests the possibility that Aurora A does not phosphorylate the GPSM2 linker in cnidarians, though it is also possible that cnidarian Aurora A has a distinct target sequence. We were unable to identify a conserved target sequence in other Aurora-A substrates, namely NEDD1, NDC80, P150-glued, MAP9, Astrin, and Haspin.

### The GPSM2 linker region is not conserved in Placozoa and Porifera

The GPSM2 linker regions in Placozoa and Porifera do not readily align with bilaterian and cnidarian sequences (Figure 4A–C). We considered whether alternate phosphorylation sites could mediate interaction with Dlg.

*T. adhaerens* (Placozoan) GPSM2 contains two predicted serine and one predicted threonine phosphorylation site between amino acids 381-420 (Supplemental Figure 3A) (Blom et al., 1999). While a previous report suggested that the threonine is the conserved phosphorylation site, we consider this possibility unlikely. Firstly, the residue is a serine in cnidarian and bilaterian animals. Secondly, we do not find sequence similarity in the flanking region. Thirdly, the proximity of this site to other phosphorylated resides is expected to present a steric challenge to Dlg, which interacts with mono-phosphorylated GPSM2 in other organisms.

*A. queenslandica* (Poriferan) GPMS2 contains two predicted serine phosphorylation sites between amino acids 328-359, presenting the same challenge (Supplemental Figure 3B). Given similarity in length and sequence to other GPSM2 homologs, we expect that the entire length of the disordered linker region is included in our reconstruction (Figure 4B). Nevertheless, we considered the possibility of potential missing exons between XP_019856151.1 and XP_003389018.1. To test for this, we reverse translated nucleotide sequences of regions with high exon character located between NW_003546593.1 (14479…19908, complement) and NW_003546593.1 (9550…10710, complement)) in BLAST (Madeira et al., 2019). We identified and translated eight potential exons and determined if these amino acid sequences contained phosphorylation sites and were similar to the bilaterian and cnidarian linkers. When determining if these exons contained phosphorylation sites, we flanked the exon amino acid sequence with the last 15 amino acids from XP_019856151.1 and the first 15 amino acids from XP_003389018.1 to ensure that the properties of the surrounding amino acids were taken into account (Supplemental Figure 3C). Only one of these (NW_003546593.1 (12781…12966, complement)) contained phosphorylation sites (Supplemental Figure 3D), but the sequence was not similar to bilaterian or cnidarian linker sequences. Our analysis suggests that Dlg and GPSM2 do not interact in Placozoa or Porifera.

## Discussion

Dlg is necessary for spindle orientation across many bilaterian models. Here we provide evidence that the interaction between Dlg and Khc73 is present throughout the animal kingdom but that the interaction between Dlg and GPSM2 evolved in Cnidaria. While it remains unclear whether or not Dlg functions as a positioning cue or to assist in microtubule capture in bilaterian spindle orientation, our results suggest that neither function is required for multicellularity.

### Dlg diverged in O. minuta

Hexactinellida is a class of sponges classified by the presence of the trabecular syncytium, a continuous multinucleated tissue that is the main component of the organism (Mehl, 1991). This class of animals is therefore an ideal system to investigate if the canonical spindle orientation machinery is necessary for nuclear-only divisions. Of the three Poriferan GUK domain sequences identified and analyzed in this study, one species: *O. minuta*, a hexactinellid, showed an unusually low average percent identity. The same trend was observed in the global Dlg sequence. Given that the GB subdomain participates in many interactions important for spindle orientation, a likely explanation for this divergence is that spindle orientation is not required for the development of the trabecular syncytium. Therefore, it would not appear necessary to orient the plane of division in cells with only nuclear, and not cytoplasmic, divisions. Furthermore, how the spindle orientation complex, consisting of many cortical proteins, could interact with nuclei towards the center of the cytoplasmic mass is unclear.

### Dlg and GPSM2 may have coevolved in C. elegans

The *C. elegans* GPSM2 orthologs, GPR1/2, demonstrate low sequence conservation with GPSM2 proteins despite maintaining its role in spindle orientation (Colombo et al., 2003; Srinivasan et al., 2003; Wavreil & Yajima, 2020). GPR1/2 share only an average of 23% identity with all other animals in our analyses. Similarly, *C. elegans* Dlg shares only an average of 43% identity with all other organisms. The observation that *C. elegans* Dlg and GPSM2 sequences share lower than expected percent identity with other bilaterians, and animals overall, is consistent with the possibility that Dlg and GPSM2 coevolved in *C. elegans.* The coevolution could be the cause or consequence of differences observed in *C. elegans* ACD (asymmetrical cell divisions) compared to other bilaterians. GPR1/2 and LIN-5 (NuMA) act downstream of cell polarity factors in *C. elegans* embryos (Colombo et al., 2003; Gotta et al., 2003; Srinivasan et al., 2003) whereas *Drosophila* Pins (GPSM2), Mud (NuMA), and polarity factors are mutually dependent on one another (Chia et al., 2008; Schober et al., 1999; Wavreil & Yajima, 2020; Yu et al., 2000). Furthermore, *C. elegans* use an evidently unique system that relies on the nematode-specific protein LET-99 to localize GPR1/2 (Krueger et al., 2010; Tsou et al., 2002). Together with previous work, our analysis suggests that spindle orientation in *C. elegans* may be best viewed as an unusual system for comparison, rather than a broadly applicable model.

### Distinction between the Cnidarian and Bilaterian linkers

While site prediction software (NetPhos) predicts phosphorylation in the cnidarian linker with > 99% confidence, the consensus sequence that we identified in these animals is markedly distinct from that in bilaterians. First, although human Aurora-A is unable to efficiently phosphorylate synthetic peptides with a charged residue at the B position, it is possible that Aurora-A can phosphorylate GPSM2 *in vivo* in cnidarians. Another possibility is that cnidarians use an alternative kinase to phosphorylate GPSM2. A strong candidate is aPKC (atypical Protein Kinase C), which is implicated in cell polarity and spindle orientation (Cui et al., 2007; Hao et al., 2010; Joberty et al., 2000; Zhu, Wen, et al., 2011)

### GPSM2 may have interphase functions in Placozoa and Porifera

If the interaction between GPSM2 and Dlg arose in Cnidaria, what is the function of GPSM2 in Placozoa and Porifera? The poriferan and placozoan GPSM2 may still be able to orient cell divisions; the GoLoco and TPR domains are conserved, indicating that the interactions between GPSM2 and NuMA and GPSM2 and Gαi are likely conserved as well. While GPSM2 may interact with a more diverse group of proteins in higher ordered organisms, it is possible that localizing NuMA to the cortex is sufficient for GPSM2 to orient spindles in Porifera and Placozoa. In the *Drosophila* wing disc epithelium, spindle orientation is not disrupted by *dlg^1P20^* mutants, which can still regulate apical-basal polarity factors but cannot interact with Pins due to a truncation in the GUK domain. Wing disc epithelium spindle orientation is also preserved in *pins^62^* mutants, which are null (Bergstralh et al., 2016). Both observed effects are due to the presence of a Pins-independent mechanism of Mud localization (Bergstralh et al., 2016; Bosveld et al., 2016; David et al., 2005).

Another possibility is that poriferan and placozoan GPSM2 is involved in interphase cell behaviors. Though interphase GPSM2 functions are less understood, GPSM2 has been shown to play a role in endothelial interphase cells. While GPSM2 is dispensable for spindle orientation in this population, it is required for endothelial cell-cell adhesion, cell-matrix adhesion, and migration (Wright et al., 2015). The disruption of cell contacts and migration in GPSM2 knock-down endothelial cells is likely due to GPSM2 regulating microtubule dynamics and focal adhesion turnover, though the exact mechanisms have not been elucidated. This finding demonstrates that GPSM2 can have functions that are independent of spindle orientation. However, a final possibility is that GPSM2 could function to both localize NuMA in metaphase cells and regulate interphase behaviors in Placozoa and Porifera.

### The interaction between Dlg and GPSM2 may be required for proper neural, but not epithelial, tissue development

Spindle orientation defects appear to have more severe consequences in nervous tissue versus epithelial tissue, as mutations in spindle orientation factors have been associated with brain abnormalities in otherwise phenotypically normal animals. Our results support the possibility that Dlg and GPSM2 interact in Cnidaria and Bilateria, but not it Porifera or Placozoa. Since the latter phyla do not have neural tissue, it is tempting to speculate that the interaction evolved to help drive neurodevelopment. However, we were unable to identify either protein in Ctenophora, which also have neurons.

Recent work in molecular phylogenetics place ctenophores as the sister group to all other animals (Moroz et al., 2014; Ryan et al., 2013). One possibility is that the common ancestor to all animal clades had neurons, but they were lost in Placozoa and Porifera. However, transcriptome analysis of ctenophores revealed that they lack, or express in non-neural tissue, the bilaterian neuronal pathway genes. These data support the idea that ctenophore nervous systems evolved independently from those in cnidarians and bilaterians (Moroz et al., 2014).

The mechanisms through which the nervous system evolved in Cnidaria and Bilateria are unresolved. A prominent hypothesis is that neurons evolved from epithelial tissue that gained the ability to sense external stimuli and transmit electrical and chemical signals to neighboring cells (Mackie, 1970). This theory is supported by the biological function of specialized cnidarian epithelial cells called nematocytes. Even in the absence of nerve cells, nematocytes can cause discharge of the nematocysts in response to chemical and mechanical stimuli (Aerne et al., 1991; Miljkovic-Licina et al., 2004). Additionally, similarities established between the epithelial septate junctions and neural synapse junctions suggest they are evolutionarily related (Banerjee et al., 2006; Harden et al., 2016) and recent work in the *Drosophila* follicular epithelium demonstrates that the epithelial cell reintegration and IgCAM-mediated axon growth occur through the same mechanism (Cammarota et al., 2020). The identification of IgCAMs in *T. adhaerens*, which contain epithelial cells but not neural, strongly suggests that the epithelial pathway was co-opted for axon growth and pathfinding when nervous tissue evolved from epithelia (Cammarota et al., 2020).

Our work may also provide insights into how epithelial cell processes may have been adopted in neurons. One possibility is that the tight regulation of asymmetric cell division (ACD) in the nervous system evolved from pre-existing ACD mechanisms. While the Insc-GPSM2 interaction is the main contributor to ACD in neuroblasts, Dlg has also been shown to regulate ACD in *Drosophila* SOP (sensory organ precursor) cells, which do not endogenously express Insc (Bellaïche et al., 2001). Additionally, Dlg is required for ACD of *Drosophila* neuroblasts in *Insc* null mutants (Siegrist & Doe, 2005). We therefore postulate that the interaction between Dlg and GPSM2 arose to regulate ACD as cell type and tissue diversity increased within the animal kingdom. Insc then co-opted this system of ACD regulation in neural tissues, where tight regulation of ACD is a crucial for normal animal development.

Several questions remain unresolved. Firstly, we were unable to identify Insc outside of the bilaterian clade, suggesting that Cnidaria do not express Insc. Does Dlg then regulate all neural ACD in this phylum? Additionally, what mechanisms regulate neural ACD in Ctenophora if their nervous system evolved independent from Cnidaria and Bilateria? The answers to these questions may further elucidate the functional consequences of interactions in the spindle orientation complex and how they relate to neurodevelopment.

The findings presented in this study indicate that the mechanisms of spindle orientation have undergone substantial evolution within the animal kingdom. While canonical spindle orientation proteins GPSM2 and Dlg can be identified in basal organisms, sequence alignments reveal that their interaction is likely not required for multicellularity. Rather, the interaction between GPSM2 and Dlg appears at the origin of neurons, suggesting it may have served as a template for asymmetrical cell division regulation in organisms with increasingly diverse and complex tissue development.

## Methods

### Protein sequence identification and alignment

The non-redundant protein sequences (nr) database and PSI-BLAST Iteration 1 (Altschul, et al., 1997) program were used for all NCBI BLAST searches (Sayers, Beck, et al., 2020; Sayers, Cavanaugh, et al., 2020) Reciprocal BLAST analysis was conducted to confirm identity. This approach confirmed that the identified sequence for *O. minuta* is not actually a different, closely related MAGUK family protein.; using *O. minuta* Dlg as the query sequence against the *H. sapiens* genome resulted in only hits against Dlg isoforms with E-values < 10^−10^.

Using *H. sapiens* GPSM2 (LGN; NP_001307967.1) and Dlg (NP_001353143.1) as the query sequence, we performed a BLAST search against common bilaterian model organisms (*M. musculus, X. laevis, D. rerio*, *S. purpuratus*, *D. melanogaster, C. gigas*). Since spindle orientation factors have not been studied in basal animals, the entire Cnidarian (taxid:6073), Placazoan (taxid:10226), Poriferan (taxid:6040) and Ctenophoran (taxid: 10197) phyla were searched for homologous sequences. With the exception of *C. elegans*, GPSM2 sequences were identified in the bilaterian models as well as *N. vectensis, A. tenebrosa, E. diaphana,* and *T. adhaerens* each with > 30% identity to *H. sapiens* GPSM2 and an E-value < 10^−10^. Similarly, Dlg sequences were identified in the bilaterian models, *N. vectensis, A. tenebrosa, E. diaphana, T. adhaerens, O. lobularis,* and *O. minuta* each with > 30% identity to *H. sapiens* Dlg and an E-value of < 10^−10^ (Supplemental Figure 1). These results indicate that the identified sequences share significant similarity with human GPSM2 (LGN) and therefore are homologs of human GPSM2 (Pearson 2013). While GPSM2 was identified in *A. queenslandica,* only a partial Dlg sequence, containing the GUK domain, was identified in *A. queenslandica* (XP_011408781.2) and therefore this organism was not included in the Dlg alignment.

Using *H. sapiens* Khc73 (KIF13B) as the query sequence (Q9NQT8), we identified potential orthologs in all organisms except for *S. purpuratus* and *T. adhaerens.* Reciprocal BLAST searching confirmed that identified sequences are the best hit Khc73 homologs in *M. musculus, X. laevis,* and *D. rerio.* However, the reciprocal BLAST for all other identified sequences indicated the best hit as KIF13A, a kinesin-motor protein that is closely related to Khc73. To confirm whether or not identified sequences are orthologs to KIF13A or Khc73, we created and analyzed the conservation domain map. While Khc73 contains an N-terminal Cap-Gly domain, KIF13A does not. Therefore, conservation at this region can be used to distinguish between the two proteins. The conservation domain map revealed a highly conserved region at the N-terminus of identified sequences, which we confirmed to be conserved at the *H. sapiens* Cap-Gly domain by analyzing the corresponding amino acid sequences. However, the identified sequence for *C. elegans* was truncated at the N-terminus and did not contain the Cap-Gly region, indicating it either is missing an exon at the N-terminus or is not the ortholog of Khc73. Therefore, *C. elegans,* along with *T. adhaerens* and *S. purpuratus,* were excluded from our analysis.

Alignments were created using COBALT (constraint-based alignment tool) (Papadopoulos & Agarwala, 2007) and were confirmed using T-coffee (Notredame et al., 2000) and ClustalX (Jeanmougin et al., 1998). Global conservation maps were created from the Multiple Sequence Alignment Viewer application where conservation was determined by relative entropy threshold of the aligned residues. Alignments from COBALT were downloaded as FASTA files and then were viewed in Jalview 2 (Waterhouse et al., 2009) where local sequence alignment figures were created.

Percent identity for matrices was calculated using EMBL-EBI Simple Phylogeny (Madeira et al., 2019). Nucleotide sequences from *A. queenslandica* exons were translated into amino acid sequences using EMBOSS EMBL-EBI Transeq (Madeira et al., 2019) with reverse set to true.

### Phosphorylation site prediction

Phosphorylation sites at predicted GPSM2 linker sequences were determined using NetPhos 3.1 (Blom et al., 1999). Serine, threonine, and tyrosine residues were considered, and graphics were modeled after the NetPhos 3.1 Server output. Only phosphorylation sites predicted with > 90% confidence were considered to be likely phosphorylation sites in this study.

### Protein structure prediction

We used the trRosetta (transformed-restrained Rosetta) algorithm to predict the structure of identified GPSM2 and Dlg sequences (Hiranuma et al., 2021; Yang et al., 2020). Identified amino acids sequences were uploaded to the Robetta server where the trRosetta algorithm was used to create a PDB file with predicted protein structures (Hiranuma et al., 2021; Yang et al., 2020). Angstrom error graphs were also obtained through the Robetta server. The PyMOL Molecular Graphics System, Version 2.4.1 was used to create 3D visualizations of protein structures. Structures were imported to PyMOL and aligned using the alignment plugin.

Predicted GPSM2 structures have conserved secondary structures at previously defined domains. Each TPR consisted of a helix-turn-helix motif while each GoLoco domain contains an alpha helix. These results provide further evidence that the sequences presented in this study are orthologous. However, due to high angstrom error estimates within unstructured regions of these proteins, including the GPSM2 linker region, these structures could not be used to predict interactions between GPSM2 and Dlg.

## Acknowledgments

We are grateful to J. David Lambert, Michael Welte, and members of the Bergstralh lab for their questions and comments.

## Author Contributions

ES designed and performed the analyses, and ES wrote the manuscript. DTB conceived and guided the project.

---

***Supplemental Figure 1. Percent Identity Matrices.*** Percent identity matrices for **A)** full length Dlg; **B)** the Dlg-GUK domain; **C)** the MAGUK-binding stalk of Khc73; and **D)** full length GPSM2.

***Supplemental Figure 2. The Cnidarian GPSM2 linker.*** Sequence alignment at the GPSM2 linker region in Cnidaria. Alignment is colored using ClustalX color scheme. Figure is adapted from Jalview 2 (Waterhouse et al., 2009).

***Supplemental Figure 3. Phosphorylation site predictions.*** Predicted phosphorylation potential of **A)** *T. adhaerens* (Placozoa) and **B)** *A. queenslandica* (Porifera) GPSM2 linker regions. Asterisks indicate the residues that are predicted to be a phosphorylation site with greater than 90% confidence. Graphs are adapted from NetPhos 3.1 (Blom et al., 1999). **C)** Reverse translated nucleotide sequences of eight regions with high exon character between NW_003546593.1 (14479…19908, complement) and NW_003546593.1 (9550…10710, complement). **D)** Of the eight potential exons identified, only one had predicted phosphorylation sites.

