## Supplemental Figure 1 for "Interaction between Discs large and GPSM2: A Comparison Across Species"

### Supplemental Figure 1 - Schiller and Bergstralh

## A

**Dlg**

| | H. sapiens | M. musculus | X. laevis | D. rerio | S. purpuratus | D. melanogaster | C. gigas | C. elegans | E. diaphana | A. tenebrosa | N. vectensis | T. adhaerens | O. lobularis | O. minuta | Average |
|---|---|---|---|---|---|---|---|---|---|---|---|---|---|---|---|
| H. sapiens | 100 | 99.59 | 87.43 | 84.33 | 59.29 | 55.34 | 59.83 | 44.57 | 59.34 | 57.23 | 58.82 | 49.5 | 54.65 | 31.9 | 61.68% |
| M. musculus | 99.59 | 100 | 87.71 | 84.05 | 59.29 | 55.34 | 59.68 | 44.57 | 59.34 | 57.23 | 58.82 | 49.36 | 54.8 | 32.06 | 61.68% |
| X. laevis | 87.43 | 87.71 | 100 | 80.35 | 60.95 | 53.39 | 57.73 | 42.98 | 59.12 | 54.87 | 56.47 | 46.57 | 52.65 | 32.42 | 59.43% |
| D. rerio | 84.33 | 84.05 | 80.35 | 100 | 59.42 | 50.85 | 56.12 | 41.41 | 58.61 | 55.39 | 55.86 | 45.05 | 50.97 | 32.43 | 58.06% |
| S. purpuratus | 59.29 | 59.29 | 60.95 | 59.42 | 100 | 55.51 | 61.55 | 44.72 | 62.3 | 62.5 | 62.39 | 48.66 | 55.32 | 33.28 | 55.78% |
| D. melanogaster | 55.34 | 55.34 | 53.39 | 50.85 | 55.51 | 100 | 63.15 | 42.81 | 57.26 | 57.28 | 55.63 | 42.56 | 52.65 | 33.42 | 51.94% |
| C. gigas | 59.83 | 59.68 | 57.73 | 56.12 | 61.55 | 63.15 | 100 | 43.97 | 62 | 61.17 | 61.71 | 46.64 | 57.72 | 33.38 | 55.74% |
| C. elegans | 44.57 | 44.57 | 42.98 | 41.41 | 44.72 | 42.81 | 43.97 | 100 | 45.34 | 42.91 | 44.5 | 35.68 | 42.23 | 30.82 | 42.04% |
| E. diaphana | 59.34 | 59.34 | 59.12 | 58.61 | 62.3 | 57.26 | 62 | 45.34 | 100 | 85.02 | 80.73 | 51.57 | 58.35 | 33.54 | 59.42% |
| A. tenebrosa | 57.23 | 57.23 | 54.87 | 55.39 | 62.5 | 57.28 | 61.17 | 42.91 | 85.02 | 100 | 79.02 | 45.94 | 58.41 | 33.75 | 57.75% |
| N. vectensis | 58.82 | 58.82 | 56.47 | 55.86 | 62.39 | 55.63 | 61.71 | 44.5 | 80.73 | 79.02 | 100 | 45.87 | 59.04 | 34.5 | 57.95% |
| T. adhaerens | 49.5 | 49.36 | 46.57 | 45.05 | 48.66 | 42.56 | 46.64 | 35.68 | 51.57 | 45.94 | 45.87 | 100 | 42.17 | 29.93 | 44.58% |
| O. lobularis | 54.65 | 54.8 | 52.65 | 50.97 | 55.32 | 52.65 | 57.72 | 42.23 | 58.35 | 58.41 | 59.04 | 42.17 | 100 | 34.33 | 51.79% |
| O. minuta | 31.9 | 32.06 | 32.42 | 32.43 | 33.28 | 33.42 | 33.38 | 30.82 | 33.54 | 33.75 | 34.5 | 29.93 | 34.33 | 100 | 32.75% |

## B

**Dlg-GUK**

| | H. sapiens | M. musculus | X. laevis | D. rerio | S. purpuratus | D. melanogaster | C. gigas | C. elegans | E. diaphana | A. tenebrosa | N. vectensis | T. adhaerens | A. queenslandica | O. lobularis | O. minuta | M. brevicollis | S. rosetta | Average |
|---|---|---|---|---|---|---|---|---|---|---|---|---|---|---|---|---|---|---|
| H. sapiens | 100 | 100 | 93.33 | 93.33 | 76.67 | 86.67 | 80 | 61.02 | 76.67 | 73.33 | 75 | 66.67 | 73.33 | 72.88 | 54.24 | 59.32 | 59.32 | 75.11% |
| M. musculus | 100 | 100 | 93.33 | 93.33 | 76.67 | 86.67 | 80 | 61.02 | 76.67 | 73.33 | 75 | 66.67 | 73.33 | 72.88 | 54.24 | 59.32 | 59.32 | 75.11% |
| X. laevis | 93.33 | 93.33 | 100 | 91.67 | 78.33 | 86.67 | 81.67 | 61.02 | 78.33 | 75 | 76.67 | 66.67 | 73.33 | 72.88 | 52.54 | 59.32 | 57.63 | 74.91% |
| D. rerio | 93.33 | 93.33 | 91.67 | 100 | 81.67 | 91.67 | 83.33 | 64.41 | 78.33 | 75 | 76.67 | 66.67 | 75 | 76.27 | 54.24 | 59.32 | 57.63 | 76.16% |
| S. purpuratus | 76.67 | 76.67 | 78.33 | 81.67 | 100 | 85 | 88.33 | 59.32 | 75 | 81.67 | 78.33 | 68.33 | 73.33 | 72.88 | 50.85 | 61.02 | 54.24 | 72.60% |
| D. melanogaster | 86.67 | 86.67 | 86.67 | 91.67 | 85 | 100 | 90 | 66.1 | 83.33 | 78.33 | 81.67 | 65 | 80 | 79.66 | 55.93 | 57.63 | 57.63 | 77.00% |
| C. gigas | 80 | 80 | 81.67 | 83.33 | 88.33 | 90 | 100 | 61.02 | 83.33 | 85 | 85 | 70 | 75 | 76.27 | 52.54 | 57.63 | 54.24 | 75.21% |
| C. elegans | 61.02 | 61.02 | 61.02 | 64.41 | 59.32 | 66.1 | 61.02 | 100 | 57.63 | 54.24 | 55.93 | 59.32 | 62.71 | 61.02 | 58.62 | 52.54 | 55.93 | 59.49% |
| E. diaphana | 76.67 | 76.67 | 78.33 | 78.33 | 75 | 83.33 | 83.33 | 57.63 | 100 | 90 | 90 | 65 | 70 | 77.97 | 54.24 | 55.93 | 50.85 | 72.71% |
| A. tenebrosa | 73.33 | 73.33 | 75 | 75 | 81.67 | 78.33 | 85 | 54.24 | 90 | 100 | 90 | 68.33 | 66.67 | 76.27 | 50.85 | 55.93 | 49.15 | 71.44% |
| N. vectensis | 75 | 75 | 76.67 | 76.67 | 78.33 | 81.67 | 85 | 55.93 | 90 | 90 | 100 | 70 | 68.33 | 72.88 | 52.54 | 57.63 | 52.54 | 72.39% |
| T. adhaerens | 66.67 | 66.67 | 66.67 | 66.67 | 68.33 | 65 | 70 | 59.32 | 65 | 68.33 | 70 | 100 | 58.33 | 67.8 | 55.93 | 55.93 | 57.63 | 64.27% |
| A. queenslandica | 73.33 | 73.33 | 73.33 | 75 | 73.33 | 80 | 75 | 62.71 | 70 | 66.67 | 68.33 | 58.33 | 100 | 69.49 | 50.85 | 55.93 | 52.54 | 67.39% |
| O. lobularis | 72.88 | 72.88 | 72.88 | 76.27 | 72.88 | 79.66 | 76.27 | 61.02 | 77.97 | 76.27 | 72.88 | 67.8 | 69.49 | 100 | 55.17 | 54.24 | 54.24 | 69.55% |
| O. minuta | 54.24 | 54.24 | 52.54 | 54.24 | 50.85 | 55.93 | 52.54 | 58.62 | 54.24 | 50.85 | 52.54 | 55.93 | 50.85 | 55.17 | 100 | 44.83 | 46.55 | 52.76% |
| M. brevicollis | 59.32 | 59.32 | 59.32 | 59.32 | 61.02 | 57.63 | 57.63 | 52.54 | 55.93 | 55.93 | 57.63 | 55.93 | 55.93 | 54.24 | 44.83 | 100 | 62.71 | 56.82% |
| S. rosetta | 59.32 | 59.32 | 57.63 | 57.63 | 54.24 | 57.63 | 54.24 | 55.93 | 50.85 | 49.15 | 52.54 | 57.63 | 52.54 | 54.24 | 46.55 | 62.71 | 100 | 55.13% |

## C

**Khc73-MBS**

| | H. sapiens | M. musculus | X. laevis | D. rerio | D. melanogaster | C. gigas | E. diaphana | A. tenebrosa | N. vectensis | A. queenslandica | Average |
|---|---|---|---|---|---|---|---|---|---|---|---|
| H. sapiens | 100 | 95.9 | 81.15 | 76.23 | 49.58 | 50.82 | 46.96 | 46.09 | 45.22 | 37.84 | 58.86% |
| M. musculus | 95.9 | 100 | 81.15 | 76.23 | 49.58 | 50 | 48.7 | 46.96 | 45.22 | 36.94 | 58.96% |
| X. laevis | 81.15 | 81.15 | 100 | 76.23 | 51.26 | 48.36 | 47.83 | 45.22 | 46.09 | 34.23 | 56.84% |
| D. rerio | 76.23 | 76.23 | 76.23 | 100 | 52.1 | 44.26 | 42.61 | 41.74 | 38.26 | 31.53 | 53.24% |
| D. melanogaster | 49.58 | 49.58 | 51.26 | 52.1 | 100 | 60.33 | 47.79 | 46.9 | 45.13 | 38.18 | 48.98% |
| C. gigas | 50.82 | 50 | 48.36 | 44.26 | 60.33 | 100 | 52.99 | 54.7 | 51.28 | 36.75 | 49.94% |
| E. diaphana | 46.96 | 48.7 | 47.83 | 42.61 | 47.79 | 52.99 | 100 | 84.55 | 81.3 | 43.75 | 55.16% |
| A. tenebrosa | 46.09 | 46.96 | 45.22 | 41.74 | 46.9 | 54.7 | 84.55 | 100 | 83.87 | 42.86 | 54.77% |
| N. vectensis | 45.22 | 45.22 | 46.09 | 38.26 | 45.13 | 51.28 | 81.3 | 83.87 | 100 | 44.64 | 53.45% |
| A. queenslandica | 37.84 | 36.94 | 34.23 | 31.53 | 38.18 | 36.75 | 43.75 | 42.86 | 44.64 | 100 | 38.52% |

Percent Identity

- 100%
- 80-99.99%
- 60-79.99%
- 40-59.99%
- 20-39.99%
- 0-19.99%

## D

**GPSM2**

| | H.sapiens | M.musculus | X.laevis | D.rerio | S.purpuratus | D.melanogaster | C.gigas | C.elegans GPR1 | C.elegans GPR2 | E.diaphana | A.tenebrosa | N.vectensis | T.adhaerens | A.queenslandica | Average |
|---|---|---|---|---|---|---|---|---|---|---|---|---|---|---|---|
| H.sapiens | 100 | 92.32 | 79.45 | 71.86 | 56.24 | 53 | 58.78 | 18.12 | 18.55 | 56.1 | 57.91 | 57.09 | 49.11 | 32.59 | 53.93% |
| M.musculus | 92.32 | 100 | 78.78 | 71.63 | 55.7 | 53.25 | 58.68 | 17.23 | 17.66 | 55.75 | 57.12 | 56.04 | 49.02 | 32.66 | 53.53% |
| X.laevis | 79.45 | 78.78 | 100 | 70.97 | 57.19 | 56.06 | 60.74 | 15.74 | 16.17 | 57.19 | 58.32 | 57.6 | 50.27 | 31.93 | 53.12% |
| D.rerio | 71.86 | 71.63 | 70.97 | 100 | 55.24 | 55.42 | 57.96 | 19.59 | 20 | 55.61 | 56.31 | 56.37 | 48.7 | 31.11 | 51.60% |
| S.purpuratus | 56.24 | 55.7 | 57.19 | 55.24 | 100 | 50.69 | 59.34 | 17.4 | 17.62 | 58.97 | 59.15 | 60.45 | 49.28 | 28.91 | 48.17% |
| D.melanogaster | 53 | 53.25 | 56.06 | 55.42 | 50.69 | 100 | 54.73 | 15.35 | 15.58 | 53.6 | 53.97 | 53.87 | 44.36 | 29.69 | 45.35% |
| C.gigas | 58.78 | 58.68 | 60.74 | 57.96 | 59.34 | 54.73 | 100 | 16.55 | 16.78 | 60.1 | 60.97 | 60.67 | 49.82 | 29.2 | 49.56% |
| C.elegans (GPR 1) | 18.12 | 17.23 | 15.74 | 19.59 | 17.4 | 15.35 | 16.55 | 100 | 96.95 | 15.42 | 15.85 | 15.84 | 17.75 | 16.15 | 22.92% |
| C.elegans (GPR 2) | 18.55 | 17.66 | 16.17 | 20 | 17.62 | 15.58 | 16.78 | 96.95 | 100 | 15.42 | 15.85 | 15.84 | 17.03 | 16.43 | 23.07% |
| E.diaphana | 56.1 | 55.75 | 57.19 | 55.61 | 58.97 | 53.6 | 60.1 | 15.42 | 15.42 | 100 | 85.81 | 82.74 | 51.83 | 28.73 | 52.10% |
| A.tenebrosa | 57.91 | 57.12 | 58.32 | 56.31 | 59.15 | 53.97 | 60.97 | 15.85 | 15.85 | 85.81 | 100 | 81.88 | 52.29 | 29.44 | 52.68% |
| N.vectensis | 57.09 | 56.04 | 57.6 | 56.37 | 60.45 | 53.87 | 60.67 | 15.84 | 15.84 | 82.74 | 81.88 | 100 | 53.89 | 29.22 | 52.42% |
| T.adhaerens | 49.11 | 49.02 | 50.27 | 48.7 | 49.28 | 44.36 | 49.82 | 17.75 | 17.03 | 51.83 | 52.29 | 53.89 | 100 | 26.61 | 43.07% |
| A.queenslandica | 32.59 | 32.66 | 31.93 | 31.11 | 28.91 | 29.69 | 29.2 | 16.15 | 16.43 | 28.73 | 29.44 | 29.22 | 26.61 | 100 | 27.90% |
