## Supplementary figures and images for "Interaction between Discs large and GPSM2: A Comparison Across Species"

### Supplemental Figure 2

#### Supplemental Figure 2 - Schiller and Bergstralh

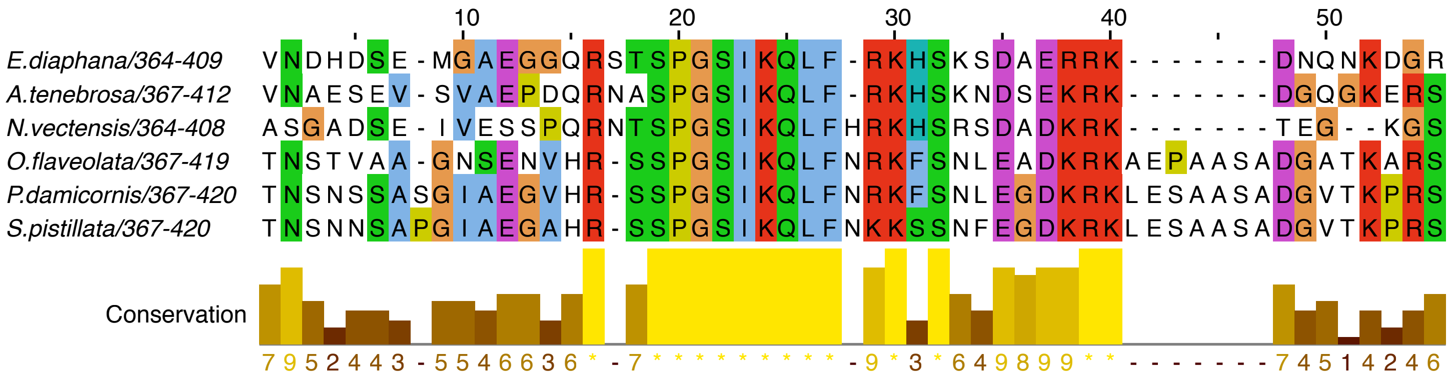
