## Supplemental Figure 3 for "Interaction between Discs large and GPSM2: A Comparison Across Species"

### Supplemental Figure 3 - Schiller and Bergstralh

**A**

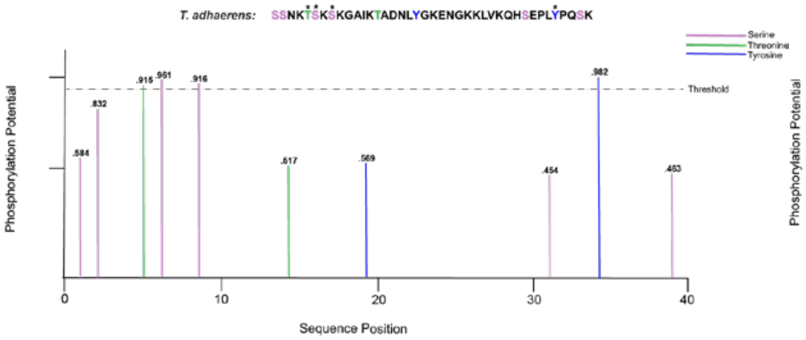


**B**

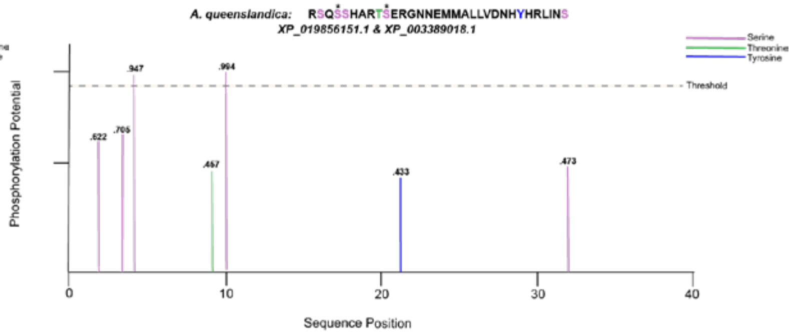


**C**

NW_003546593.1 Exons

| complement | |
|---|---|
| 10711...10795 | MAETGISVHS RMMAQDIFRI LSIAFIKEX |
| 10901...11001 | LVCFKTRQPL ACSVLNGLTE PSANVSLSY |
| 12193...12217 | IMRTPLGYLY NKDVCPNWYK PI |
| 12593...12642 | RDYSLRKTAN GGYYQYX |
| 12781...12966 | PFVKLRKXMS VCGPRIIRCR EYXHSKHCMG SMDTPTSYKT SKLTQTQLAS QPQLTHQXXL EL |
| 13327...13360 | SCGREVRRT |
| 13424...13473 | GTSIFRKKSI PLHVT |
| 14350...14412 | FCPVLIYLNK FLSSTLDVLF Y |

**D**

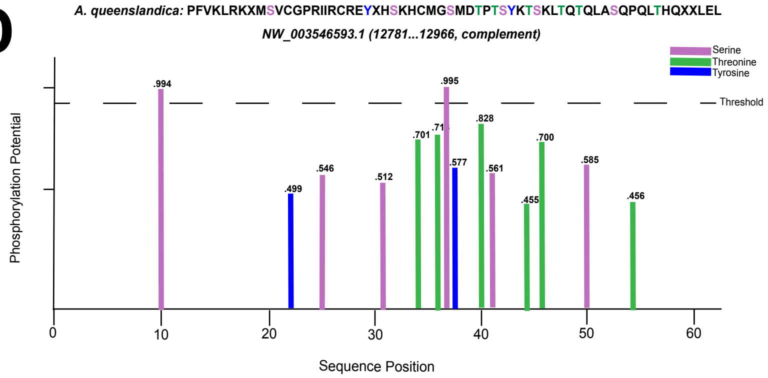
